# Multi-step engineered adeno-associated virus enables whole-brain mRNA delivery

**DOI:** 10.1101/2024.06.04.597261

**Authors:** Weiya Bai, Dong Yang, Yaqing Zhao, Guoling Li, Zhen Liu, Panzhi Xiong, Haifeng Quan, Xiaoqing Wu, Peng Chen, Xiangfeng Kong, Xuchen Wang, Hainan Zhang, Yingsi Zhou, Tong Li, Yuan Yuan, Xuan Yao, Linyu Shi, Hui Yang

## Abstract

Adeno-associated viruses (AAVs) are commonly used vectors for DNA delivery in gene therapy. Here we developed a system that enables the AAV shell to package mRNAs by multi-step introduction of RNA-packaging components and modification of AAV Rep proteins. The resultant mRNA-carrying AAVs (RAAVs) retained most properties of conventional AAVs, including capsid composition, virus morphology, and tissue tropism. These RAAVs could mediate mRNA transfer into target cells and tissues, leading to transient expression of the functional protein. Importantly, intravenously injected RAAVs efficiently crossed the blood-brain barrier (BBB) and infected the whole mouse brain. Thus, the DNA viral vector could be modified for RNA delivery, and our RAAV represents the first highly efficient BBB-crossing mRNA delivery system that could be used for therapeutic purposes via whole-brain infection.

## Main Text

Messenger RNAs (mRNAs) have emerged as a new category of therapeutic agents for prevention and treatment of many diseases. For introducing exogenous mRNAs *in vivo*, the delivery system needs to protect the nucleic acid from degradation and allow effective cellular uptake and mRNA release^1^. Lipid nanoparticles (LNPs) have been developed as a RNA-delivery system, and used clinically for the delivery of siRNA drugs^2^ and mRNA vaccines^3-5^, as exemplified by its use in delivering antigen mRNAs as coronavirus disease 2019 (COVID-19) vaccines^3-5^. In addition, virus-like particles (VLPs) are used as mRNA-delivery tools to combine the high infection efficiency of viral vectors and the transient nature of introduced mRNA^6-10^. However, systemic injection of mRNA-delivering LNPs and VLPs was found to target mainly liver^11^, with low efficiency for delivery into many\ other tissues, particularly the central nervous system (CNS) due to the presence of blood-brain barrier (BBB). The use of naturally occurring and newly engineered AAV capsids is a promising strategy for targeting non-liver tissues, such as CNS^12^, skeletal muscle^13^, and heart^14^. AAV is a small, non-enveloped virus that could package a single-stranded DNA (ssDNA)^15^ and has been engineered for DNA delivery, by replacing all viral protein-coding sequences with the therapeutic gene expression cassette between two required packaging signals (inverted terminal repeat, ITR)^16^. Unlike retroviruses-derived VLPs in which virus assembly and genome encapsidation occur simultaneously, synthesized AAV genome is pumped into a pre-assembled capsid in a 3’ to 5’ direction via the viral DNA helicase/ATPase activity of Rep proteins (Rep78, Rep68, Rep52, and Rep40) as the motor^17,18^. Non-structural Rep78/68 proteins also serve as the ‘bridge’ between the ssDNA genome and the pre-assembled AAV capsid during virus packaging (Fig. 1a)^15,18^. Based on these findings, we hypothesized that replacing ITR with RNA packaging signals (RPSs) and enabling the binding of Rep78/68 proteins to RPS-harboring mRNAs may convert the AAV into an RNA-packaging virus.

**Fig. 1.**
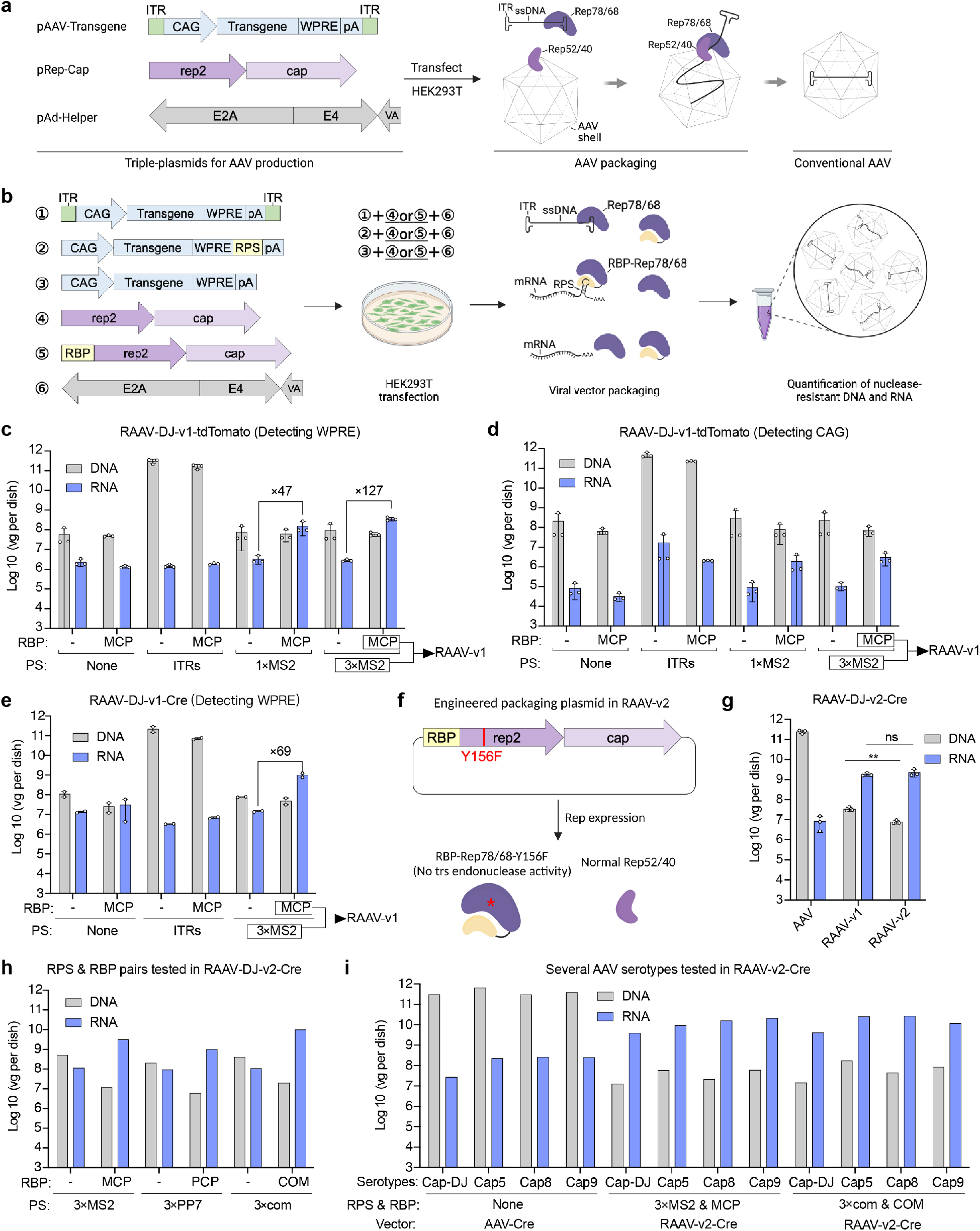
Establishment of RAAV systems for producing mRNA-carrying AAVs. **a**, The principle of conventional AAV production. ITRs, inverted terminal repeats; CAG, CAG promoter; WPRE, Woodchuck Hepatitis Virus Posttranscriptional Regulatory Element; pA, poly(A). **b**, Schematic of the establishment of the RAAV system. RPS, RNA packaging signal; RBP, RPS binding protein. The RBP is fused to the N terminus of Rep78/68. **c**,**d**, Quantification of the tdTomato coding RNA and DNA encapsidated in RAAVs and AAVs by RT-qPCR and qPCR using primers targeting WPRE (**c**) and CAG promoter (**d**). RAAV-v1, the first generation of RAAV system; MCP, MS2 coat protein; 1× or 3×MS2, one copy or three copies of MS2 stem-loops; vg per dish, vector genomes per 15-cm dish. Data are shown as individual data points and mean values ± SD for n=3 biological replicates. **e**, Quantification of the Cre coding RNA and DNA encapsidated in RAAVs and AAVs by RT-qPCR and qPCR using primers targeting WPRE. Data are shown as individual data points and mean values ± range for n=2 biological replicates. **f**, The principle of the RAAV-v2 system. The Y156F mutation was introduced into the MCP-fused Rep78/68 in RAAV-v1 to abolish their endonuclease activities, thus hindering undesired DNA release and packaging. **g**, Quantification of the Cre RNA and DNA encapsidated in RAAVs generated by RAAV-v2 system by RT-qPCR and qPCR using primers targeting WPRE. Data are shown as individual data points and mean values ± SD for n=3 biological replicates; unpaired, two-tailed t-test; **P < 0.01; ns, not significant. **h**, Two other RPS/RBP pairs were tested in the RAAV-v2 system. PP7, PP7 binding site; PCP, PP7 bacteriophage coat protein; com, Com binding site; COM, phage COM protein. **i**, Different AAV serotypes were tested in the RAAV-v2 system.

## Multi-step AAV engineering for mRNA-carrying capability

The AAV is conventionally produced by co-transfection of transgene plasmid, packaging plasmid, and helper plasmid (pAd-Helper). In the first step of our new AAV design, we removed both ITRs from the transgene plasmid (expressing tdTomato) and introduced RPSs using either one or three copies of MS2 stem-loops (1× or 3×MS2) at the 3’ end of the transgene cassette (between WPRE and poly-A tail). In parallel, a transgene plasmid with no ITR and MS2 was constructed as a negative control (Fig. 1b). For the packaging plasmid, we fused MS2 coat protein (MCP) to the N-terminus of Rep78/68 to enable its binding to MS2 in the specific mRNA transcribed from the transgene plasmid. Another packaging plasmid with no MCP was used as a negative control (Fig. 1b). The MS2/MCP pair comes from the MS2 bacteriophage, with the MCP specifically binding to the MS2 RNA stem-loops in the phage’s genome^19^. Here, MS2 and MCP were chosen based on the previous finding that this bacteriophage-derived MCP interacts with MS2 to direct the packaging of mRNA cargoes into modified VLP particles^20-22^. We co-transfected HEK293T cells with the new transgene plasmid and packaging plasmid in the presence of the pAd-Helper to produce the mRNA-carrying AAVs (termed “RAAVs”) (Fig. 1b). We also generated conventional AAVs for comparison (Fig. 1a).

After harvesting RAAV particles from the producer cells and supernatants, we analyzed the packaged nucleic acid to determine whether the AAV capsid contained the MS2-containing mRNA. To avoid the high background plasmid signals, we treated the virus stock with nucleases before extracting AAV capsid-protected nucleic acids. The extracted nuclease-resistant RNA and DNA were quantified by RT-qPCR and qPCR. Besides, two pairs of qPCR primers were designed to distinguish packaged DNAs and RNAs. Specifically, CAG-targeting primers were used for detecting DNA only, while WPRE-targeting primers detected both DNA and RNA. As expected, packaging of DNA was essentially eliminated after removing ITRs, since the detected DNA titer was about 4 orders of magnitude lower than that found in the conventional AAV (Fig. 1c,d). In the meantime, due to the introduction of MCP-fused Rep78/68 into the packaging system after removing ITRs, mRNA-containing RPSs with 1× or 3×MS2 were efficiently packaged into AAV capsids. This resulted in mRNA titers for ‘1×MS2’ and ‘3×MS2’ groups 47- and 127-fold of that found for the no MCP group, respectively. We thus used the 3×MS2 & MCP combination for the first generation RAAV system (termed “RAAV-v1”) (Fig. 1c,d). To confirm that the RAAV-v1 is effective for packaging other mRNA transgenes, we demonstrated that Cre mRNA could also be packaged with a titer similar to that of tdTomato mRNA (Fig. 1e).

Although most of the packaged nucleic acids in the above RAAVs were found to be mRNA, we also detected a small number of DNAs (4.8% of the packaged nucleic acids) (Fig. 1e). We reasoned that these DNAs in RAAV vectors could come from non-specific recognition and cleavage of the plasmid by the endonuclease activity of Rep78/68 proteins^15^. We thus constructed the “RAAV-v2” system by introducing the ‘Y156F’ mutation to the MCP-fused Rep78/68 protein (MCP-Rep78/68^Y156F^) in the RAAV-v1 to abolish their endonuclease activity but retain the helicase/ATPase activity (Fig. 1f)^23^. RAAV-v2 indeed showed significantly reduced DNA packaging without affecting the RNA packaging, as compared to that found in RAAV-v1 (Fig. 1g).

To demonstrate that the approach of using MS2/MCP in the AAV engineering for obtaining RAAVs could be generalized, we examined the mRNA-packaging efficiency of the RAAV-v2 system using two other pairs of RPS/RBP : (1) PP7 binding site and PP7 bacteriophage coat protein (PP7/PCP) and (2) Com binding site and phage COM protein (com/COM)^24,25^. Transgene plasmids harboring three copies of RPS (3×PP7 and 3×com) and their corresponding packaging plasmids (containing PCP- or COM-fused Rep78/68^Y156F^) were constructed. RAAVs were produced, purified, and titrated as described above. We found that both PP7/PCP and com/COM pairs indeed conferred markedly elevated mRNA packaging capability of RAAV, compared to that found for the conventional AAV (Fig. 1h). We also found that the mRNA packaging efficiency of RAAVs (with MS2/MCP or com/COM pair) was slightly higher when AAV capsids (capsid 5, 8, and 9) other than capsid DJ were used (Fig. 1i).

## Helicase engineering and cargo sequence optimization

Although the above engineering procedure substantially improved specific mRNA packaging of RAAV and reduced undesired DNA packaging, the titer of mRNA in RAAV still needs to be improved. The Rep proteins of AAV contained a helicase/ATPase (helicase for short) for DNA packaging (Fig. 2a)^26^. Although the above results showed that this helicase could also translocate mRNA, we surmised that the mRNA packaging efficiency may be elevated by engineering helicase via mutagenesis. The AAV helicases belong to the superfamily 3 (SF3) helicases, which contain four conserved motifs, A, B, B’, and C that constitute the core of the helicase active site^26,27^. A conserved arginine finger is located after motif C (Fig. 2a). SF3 helicases could be encoded by both DNA and RNA viruses^28,29^. We downloaded 98 SF3 helicase-containing viral protein sequences (22 ssDNA and 76 ssRNA) from Genbank and Uniprot. The core sequences of these 98 helicases were aligned and phylogenetically analyzed via AlignX (Fig. 2b). They were also aligned by MUSCLE, and full sequences of randomly selected 23 helicase-containing viral proteins were aligned via AlignX. These analyses revealed multiple highly conserved regions across all viral helicases, as well as some divergent loci between ssDNA and ssRNA viruses. We focused on the divergent loci within conserved regions and identified 114 amino acids as candidates for mutagenesis. We conducted single-codon substitution in the helicase of the RAAV-v2 and identified 13 of 114 mutated helicases that exhibited over 3-fold enhancement in mRNA packaging. Subsequent motif-directed combinatorial mutagenesis for multiple (2 to 5) amino acid substitutions, based mainly among the mutagenesis site for the above 13 helicases, resulted in 68 helicases with multiple mutations at the same or different motifs. Among the 68 mutated helicases, 6 yielded mRNA packaging capability at least 4.1-fold of that found for wild-type helicase. The combined mutation 4 achieved the highest (12.7-fold) efficiency (Fig. 2c,d), and mutated helicase was used as the “RAAV-v3” system in following experiment. Furthermore, the DNA titer of this RAAV-v3 was 5-fold lower than that of RAAV-v2 (Fig. 2c). Overall, the RAAV-v3 system now achieved ∼6,000-fold higher mRNA packaging and ∼14,100-fold lower DNA packaging, as compared with those of conventional AAV. Consequently, the mRNA titer in RAAV-v3 was only 8.93-fold lower than the DNA titer in AAV (Fig. 2d).

**Fig. 2.**
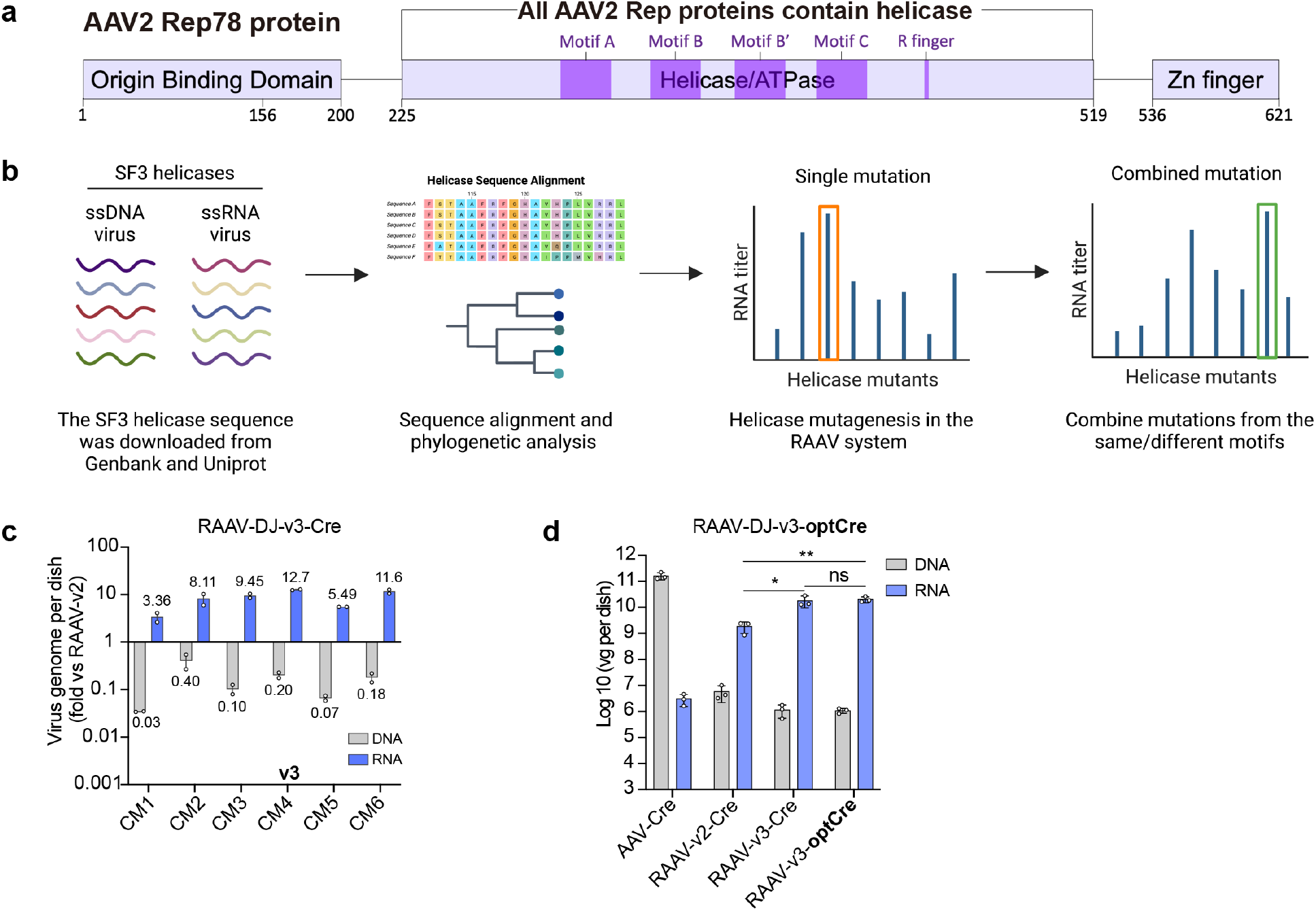
Helicase engineering to improve RAAV productivity. **a**, Schematic of the helicase domain in AAV2 Rep78 protein. All AAV2 Rep proteins (including Rep78, Rep68, Rep50, and Rep42) contain the helicase domain. **b**, Workflow for the helicase mutagenesis experiment. An artificial capsid DJ was used to produce RAAVs. ssDNA, single-stranded DNA; ssRNA, single-stranded RNA. **c**, Compare the RNA and DNA packaging capability of mutated helicases to the wild-type helicase in the RAAV-v2 system. Data are shown as individual data points and mean values ± range for n=2 biological replicates. CM, combined mutation. **d**, Quantification of the Cre RNA and Cre DNA encapsidated in RAAVs generated by RAAV-v3 system by RT-qPCR and qPCR using primers targeting WPRE. optCre, optimized Cre coding sequence 4. vg per dish, vector genomes per 15-cm dish. Data are shown as individual data points and mean values ± SD for n=3 biological replicates; unpaired, two-tailed t-test, *P < 0.05, **P < 0.01; ns, not significant.

In addition to its capability of selective mRNA packaging, the efficiency of expressing exogenous proteins in cells by our RAAVs also depends on the translation of the delivered mRNA. Thus, we further pursued codon optimization of mRNA to improve its protein translation. The Cre-coding sequence was optimized using two codon optimization online tools. We introduced RAAV (with capsid DJ) carrying optimized Cre-coding sequences (RAAV-Cre opt1 to opt4) into cultured mouse embryonic fibroblasts (MEFs) isolated from Cre reporter mice (Ai9 mice harboring *loxP*-tdTomato). The RAAV carrying original Cre-coding sequences (RAAV-Cre) was used for comparison. Five days after infection, we found that RAAV-Cre opt4 yielded significantly higher percentages of tdTomato^+^ cells than RAAV-Cre at the same multiplicity of infection (MOI, calculated as vector genomes per cell) without affecting the mRNA titer (Fig. S1), indicating elevated Cre protein expression. Therefore, RAAV-v3 with Cre mRNA using opt4 (RAAV-v3-optCre), generated by the combination of helicase mutagenesis and Cre coding sequence optimization, was currently the most efficient RAAV vector (Fig. 2d) and was used in all following experiments, except indicated otherwise.

## Further characterization of RAAVs

A comprehensive assessment of the properties of RAAV-v3-optCre was further conducted, using silver staining of protein composition of the RAAV with the SDS-PAGE method and visualization of RAAV particles with transmission electron microscopy. We found that RAAVs and AAVs were indistinguishable in the capsid composition and morphology (Fig. 3a,b). To assess the specificity and integrity of the genome of RAAV, we extracted and analyzed the genomes of RAAV-v3-optCre and AAV-Cre on denaturing agarose gels stained with SYBR™ Green II. We observed a 2000∼2400 nt band (consistent with the expected size of Cre mRNA in RAAV) that was resistant to DNase I but not RNase I, indicating that most packaged genomes in RAAV were intact mRNAs (Fig. 3c,d and Fig. S2), whereas the 3265 nt-long ssDNA in AAV-Cre was highly susceptible to degradation by DNase I but not RNase I. In addition, mRNA sequencing on whole-cell lysate and virus-like particle (VLP) fraction (after removing residual unencapsidated RNA by nuclease) was performed to identify various RNA species in the VLP fraction (Fig. 3e), with and without the presence of MCP. We found that MCP markedly and specifically elevated the amount of full-length optCre transcripts in the VLP fraction, but only slightly increased the optCre transcripts in whole-cell lysates (Fig. 3f,g and Fig. S3).

**Fig. 3.**
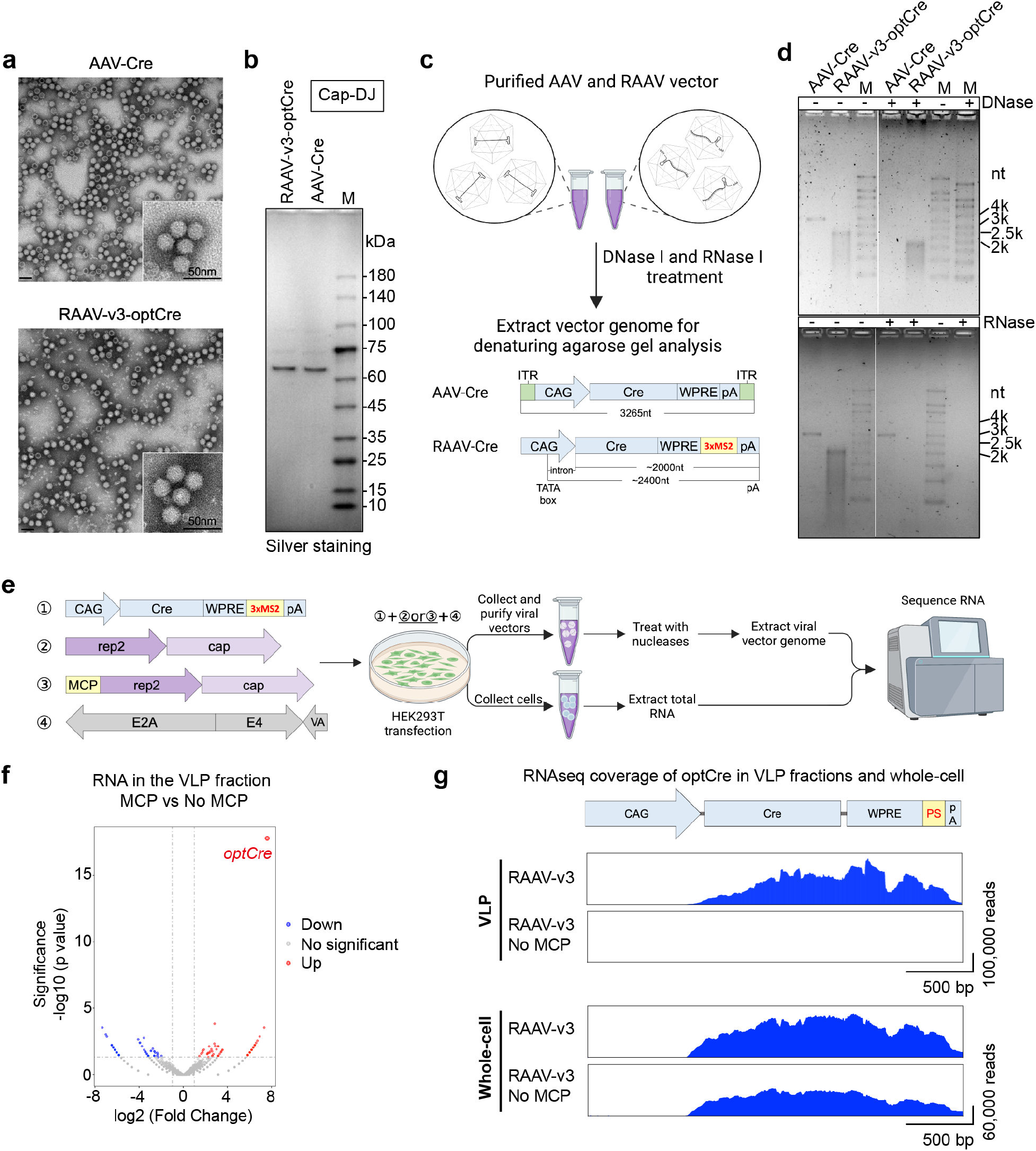
Characterization of the properties of RAAVs. **a**, Morphology analysis of RAAV and AAV by transmission electron microscopy. An artificial capsid DJ was used to produce RAAV and AAV. The bottom images are magnified versions of the image. Scale bars, 50 nm. **b**, Analyzing the composition of AAV and RAAV by silver staining. **c**, Schematic of viral vector genome analysis on denaturing agarose gel. The theoretical size of the genome of RAAV-DJ-v3-optCre and AAV-DJ-Cre are displayed in the image. **d**, Analyzing the genome of RAAVs and AAVs on a denaturing agarose gel stained with SYBR™ Green II. DNaseI and RNaseI treatment groups were set to identify the RAAV genome. M, RNA marker. **e**, Workflow for sequencing-based analysis of RAAV genome. An artificial capsid DJ was used to produce RAAVs and AAVs. **f**, Differential mRNA abundance and significance of the VLP fraction in the presence or absence of MCP. **g**, Alignment of sequencing reads showing sequencing coverage of the optCre mRNA from (**e**,**f**).

## Transient protein expression of RAAV-delivered mRNA in cells

Next, we explored the cell infectivity of RAAVs in parallel with conventional AAVs by the Cre reporter system (*loxP*-tdTomato), and the vector genome (vg) titer was used for the MOI calculation. RAAV-Cre and conventional AAV-Cre vectors were applied for 12 hours to cultured Ai9-MEFs, and the percentage of tdTomato^+^ cells was analyzed by flow cytometry 5 days after infection (Fig. 4a). The results showed that RAAV-v2-Cre, RAAV-v3-Cre, and RAAV-v3-optCre all achieved the successful transfer of Cre mRNA and expression of functional Cre protein, as indicated by tdTomato fluorescence, and the infection ability of RAAV-v3-optCre was comparable to that of AAV-Cre (Fig. 4b and Fig. S4). By contrast, negative controls of RAAV-v3-optCre (no MS2, no MCP, or no Cap) were also prepared and none showed obvious infectivity (Fig. 4b). For the infection assay, the DNA amount was normalized among RAAV, RAAV with no MS2, and RAAV with no MCP, and the infection volume was normalized between RAAV and RAAV with no Cap. Since RAAV negative controls contained significantly lower amounts of mRNA than RAAV, the infectivity of RAAV could be attributed to encapsidated mRNA, rather than residual DNA.

**Fig. 4.**
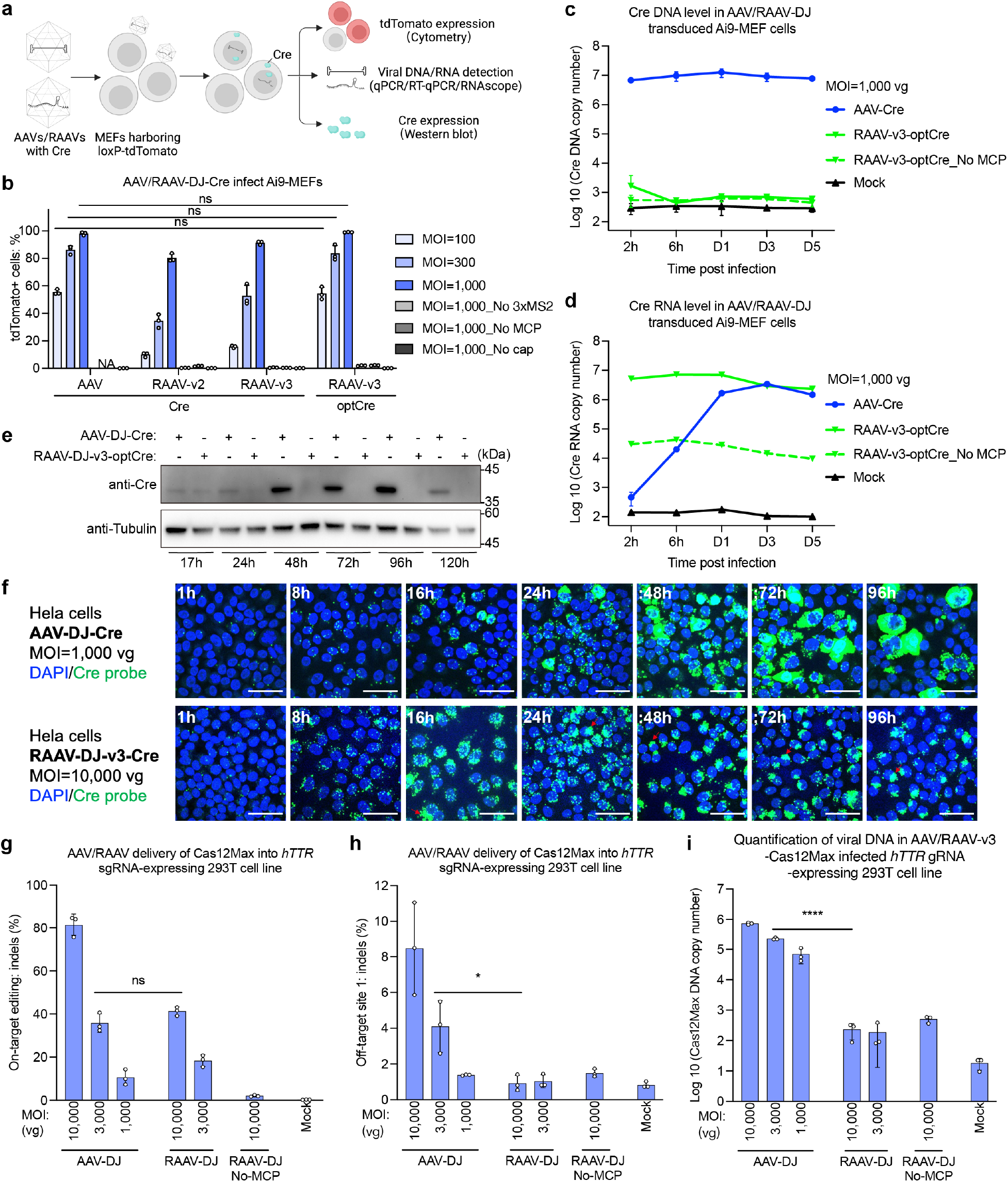
RAAVs enable efficient transfer of mRNA to target cells to transiently express functional proteins. **a**, Schematic of RAAV *in vitro* infection assay. An artificial capsid DJ was used to produce RAAVs, AAVs, and negative controls of AAVs/RAAVs (no MS2, no MCP and no Cap). Vector genome titer was used for MOI calculation. DNA amounts were normalized among RAAV, RAAV with no MS2, and RAAV with no MCP. The volume was normalized between RAAV and RAAV with no Cap for infection. For all Ai9-MEFs infection experiments, cells were plated on 48-well plates at a density of 5E4 cells per well 24 hours before infection. **b**, Investigating the infectivity of RAAVs by analyzing the percentage of tdTomato^+^ Ai9-MEFs 5 days after infection by cytometry. Data are shown as individual data points and mean values ± SD for n = 3 biological replicates, multiple unpaired t-test; ns, not significant. **c**,**d**, Time course of Cre DNA (**c**) and mRNA (**d**) levels in RAAVs/AAVs infected Ai9-MEFs. Mock: uninfected control. Data are shown as mean values ± SD for n = 3 biological replicates. **e**, Western blot analysis of Cre protein amount and lifespan in RAAV/AAV infected Ai9-MEFs. RAAV-DJ-v3-optCre infected Ai9-MEFs at MOI of 3,000 vg, and AAV-DJ-Cre infected Ai9-MEFs at MOI of 300 vg. Tubulin is used as a loading control. **f**, Spatial distribution of viral RNA and DNA in RAAV/AAV infected cells over time. Hela cells were infected with RAAV/AAV. At various time points after infection, the cells were fixed and processed for RNAscope analysis. Nuclei were visualized using DAPI staining. Viral RNA and DNA were detected with a DNA probe that binds to the Cre mRNA and DNA. Red arrow, aggregated RNA signal in the cytoplasm; Scale bars, 50 μm. **g**, Indels at the *hTTR* locus in *hTTR-*gRNA-HEK293T cells infected with RAAV-DJ-v3- or AAV-DJ-Cas12Max. Indels were quantified by NGS 120 hours after viral vector addition. **h**, NGS analysis of the off-target effect in *hTTR*-gRNA-HEK293T cells infected with RAAV-DJ-v3- or AAV-DJ-Cas12Max. The off-target site for *hTTR*-gRNA was predicted by Cas-OFFinder^***34***^. Indels at the predicted off-target site were quantified by NGS 120 hours after RAAV/AAV infection. **i**, Quantification of Cas12Max DNA copy number in RAAV/AAV infected *hTTR-*gRNA-HEK293T cells. (**g-i**) Mock: uninfected control. Vector genome titer was used for MOI calculation. Data are mean values ± SD with n = 3 biological replicates, unpaired, two-tailed t-test; *P < 0.05, ****P < 0.0001; ns, not significant.

To determine the exact level and lifespan of the viral vector-derived mRNA and the translated Cre recombinase, we infected cultured Ai9-MEFs with RAAV-v3-optCre, and the cells were collected at various time points for assaying the amounts of Cre DNA, mRNA and protein (Fig. 4a). We found that the Cre DNA level in cells infected with RAAV with and without MCP was the same as the background level found in uninfected control (MOCK) cells (Fig. 4c). The RAAV-derived mRNA could be detected in these cells as early as 2 hours post-infection, peaked at 6 hours, and decreased afterward to a lower level of ∼ 30% of the peak level by Day 5 (Fig. 4d). By contrast, the amount of Cre mRNA in AAV-Cre-infected cells was extremely low (∼ 0.01% of those in the RAAV group) at 2 hours post-infection, and then markedly elevated by 7,400-fold at day 3 (Fig. 4d). As expected, the transcription inhibitor actinomycin D largely abolished the mRNA elevation by AAV infection but had no effect on the mRNA elevation by RAAV infection (Fig. S5). Furthermore, we found that both infections did not affect the expression of housekeeping mRNA for *mGAPDH* and DNA for *36B4*^30^, confirming specific infection and gene expression induced by the viral vectors (Fig. S6). At the protein level, unlike AAV-Cre which expressed Cre persistently after infection, the expression of Cre in RAAV-infected cells could be detected as early as 17 hours post-infection and disappeared after one day (Fig. 4e). To express an equivalent amount of Cre at 17 hours post-infection, the viral genome copy number for RAAV (MOI=3,000 vg) was found to be about 10-fold of that for AAV (MOI=300 vg) (Fig. 4e), consistent with our expectation that infected AAV-Cre could transcribe multiple copies of Cre mRNAs, leading to higher amounts of protein expressed. Furthermore, the RNAscope assay showed that RAAV entered the nucleus as early as 8 hours post-infection in Hela cells like conventional AAV^31^ (Fig. 4f), and the nuclear entry of RAAV could be inhibited by bafilomycin A1 that interfered with viral infection^32^ but not by actinomycin D that only affected mRNA transcription induced by AAV (Fig. S7). Notably, while the qPCR assay showed mRNAs remained detectable 5 days after RAAV infection, the expressed Cre protein disappeared by 1 day, suggesting termination of the expressed mRNAs (Fig. 4d,e). This is in line with the finding that although mRNA exit from the nucleus was observed as early as 8 hours post-infection, with increasing cytoplasmic aggregation of mRNAs afterward (possibly involving mRNA sequestration within P-bodies), a process known to downregulate translation^33^ (Fig. 4f). Taken together, our findings showed that RAAV could mediate successful mRNA transfer to cultured cells and transiently express functional proteins, via trafficking to the nucleus similar to the conventional AAV. Thus, RAAV could also be used for introducing exogenous RNAs that regulate nuclear RNAs in the cell.

To expand the applicability of the RAAV system, we examined whether RAAV-v3 could mediate the functional transfer of a large CRISPR-Cas12Max transcript (3774 nt), which exhibits editing activity comparable to Cas9^34^. We transduced RAAV-v3-Cas12Max into 293T cells that constitutively express a guide RNA (gRNA) targeting *hTTR*^*34*^ and analyzed gene editing efficacy 5 days after infection. The results showed that RAAVs were able to functionally transfer Cas12Max mRNAs, leading to 41.3 ± 1.9 % insertions and deletions (indels) at a MOI of 10,000 vg in recipient cells, with background-level editing at predicted off-target sites as that found in untreated cultured cells (Fig. 4g,h). Furthermore, at a similar level of on-target editing efficiency, the editing rate at the off-target site 1 of RAAV-Cas12Max (at MOI of 10,000 vg group) was 4.4- fold lower than that of AAV-Cas12Max (at MOI of 3,000 vg) (Fig. 4g,h). Finally, only trace amounts of viral vector DNA were detected in RAAV-Cas12Max-infected cells (Fig. 4i), greatly reducing the possibility of DNA insertional mutations^35^. Overall, these results indicated that RAAV-mediated mRNA delivery provides a gene editing method with reduced off-target effects.

## RAAV delivery of mRNAs exhibits tissue tropism

To assess the *in vivo* delivery efficacy of RAAVs, we injected RAAV-v3-Cre, RAAV-v3-optCre, and AAV-Cre with capsid DJ into the hippocampus of adult Ai9 mice. Four weeks after injection, we analyzed the hippocampal expression of tdTomato and Cre (Fig. S8a). The average percentage of tdTomato+ cells in hippocampi injected with RAAV-v3-Cre (24.2 ± 7.1 %, n = 3) and RAAV-v3-optCre (29.9 ± 6.0 %) was lower than that observed in hippocampi injected with AAV-Cre, a DNA vector, at the same genome dose (1E8 vg/hippocampus) (60.2 ± 24.0 %), but was comparable to that of hippocampi injected with a lower dose (1E7 vg/hippocampus) of AAV-Cre (30.8 ± 4.6 %) (Fig. S8b,c). In addition, Cre expression could only be detected in hippocampi infected with AAV-Cre (Fig. S8b,c), whereas no viral vector-associated Cre proteins was detected in RAAV-Cre-infection hippocampi (Fig. S8). In comparison, no tdTomato^+^ or Cre^+^ cell was observed in the control mice treated with RAAV with no MCP (Fig. S8b,c). Thus, RAAVs could efficiently deliver Cre mRNAs into the mouse hippocampus, and transiently express functional Cre protein at a safe dose.

The AAV capsids isolated from various mammals^36^ or engineered artificially^31^ exhibit a wide variety of cell and tissue tropisms. Since the capsid is a major determinant of cell/tissue tropism of AAVs^36^, we next examined whether RAAV vectors could retain their original capsid tropism. Intravenously injected AAV with the capsid PHP.eB is known to exhibit higher infection efficiency in the brain and lower infection in the liver compared to AAV with capsid 9^12,37,38^. We generated Cre-coding RAAVs and AAVs with either capsid PHP.eB or capsid 9 and intravenously injected each viral vector into the Cre reporter mouse Ai9. The mice were sacrificed 4 weeks after infection to analyze tdTomato and Cre expression in the liver and brain tissues (Fig. 5a). For capsid 9, we found high percentages of tdTomato^+^ cells (74.7%) and Cre^+^ cells (41.3%) in the liver of AAV9-Cre injected mice (at a dose of 1E11 vg). In comparison, the percentage of tdTomato^+^ cells in the liver of RAAV9-v3-optCre-injected mice was increased from 4.8% to 74%, as the vector dose increased from 1E11 vg to 1E12 vg, while no Cre^+^ cells were detected in the liver of RAAV9-v3-optCre infected mice as expected (Fig. 5b-d). In addition, both AAV9-Cre and RAAV9-v3-optCre yielded no obvious infection in the brain (Fig. 5e). As for the capsid PHP.eB, we found much lower liver infection in AAV-PHP.eB-Cre and RAAV-PHP.eB-Cre infected mice compared with that of AAV9-Cre and RAAV9-Cre infected mice at the same dose, respectively (Fig. 5b). In contrast, we found clear infection tropism for brain tissues, with high percentages of tdTomato^+^ cells in various brain tissues of AAV-PHP.eB-Cre-infected mice (12.6%, cortex; 6.4%, hippocampus; 21.4%, thalamus; 11.3%, striatum; 20.9%, midbrain), as compared to those found in AAV9-Cre-injected mice (at a dose of 1E11 vg) (Fig. 5e-g), consistent with previous studies^12,37,38^. Interestingly, we found that RAAV-PHP.eB-v3-optCre infected mice also effectively infected the whole brain, showing high percentage of tdTomato^+^ cells (36.1%, cortex; 9.4%, hippocampus; 43.4%, thalamus; 37.6%, striatum; 38.9%, midbrain) at a dose of 1E12 vg (Fig. 5e-g), indicating that RAAV-PHP.eB shares the same tropism with AAV-PHP.eB. However, due to the short lifetime of RAAV-PHP.eB-delivered mRNA and translated Cre, we only observed Cre expression in AAV-PHP.eB-infected brains but not in RAAV-PHP.eB-infected brains (Fig. 5h). As controls, very few tdTomato^+^ cells were observed in the control mice treated with RAAV9 or RAAV-PHP.eB with no MCP (Fig. S9). In addition to the tissue tropism, RAAVs also retained their cellular tropisms, as shown by results using RAAV with various capsids to infect different types of culture cells (Fig. S10). Overall, these results indicated that RAAVs retained the infection tropism from their capsids and could mediate specific infection to target cells, tissues, and organs to transiently express functional proteins of interest.

**Fig. 5.**
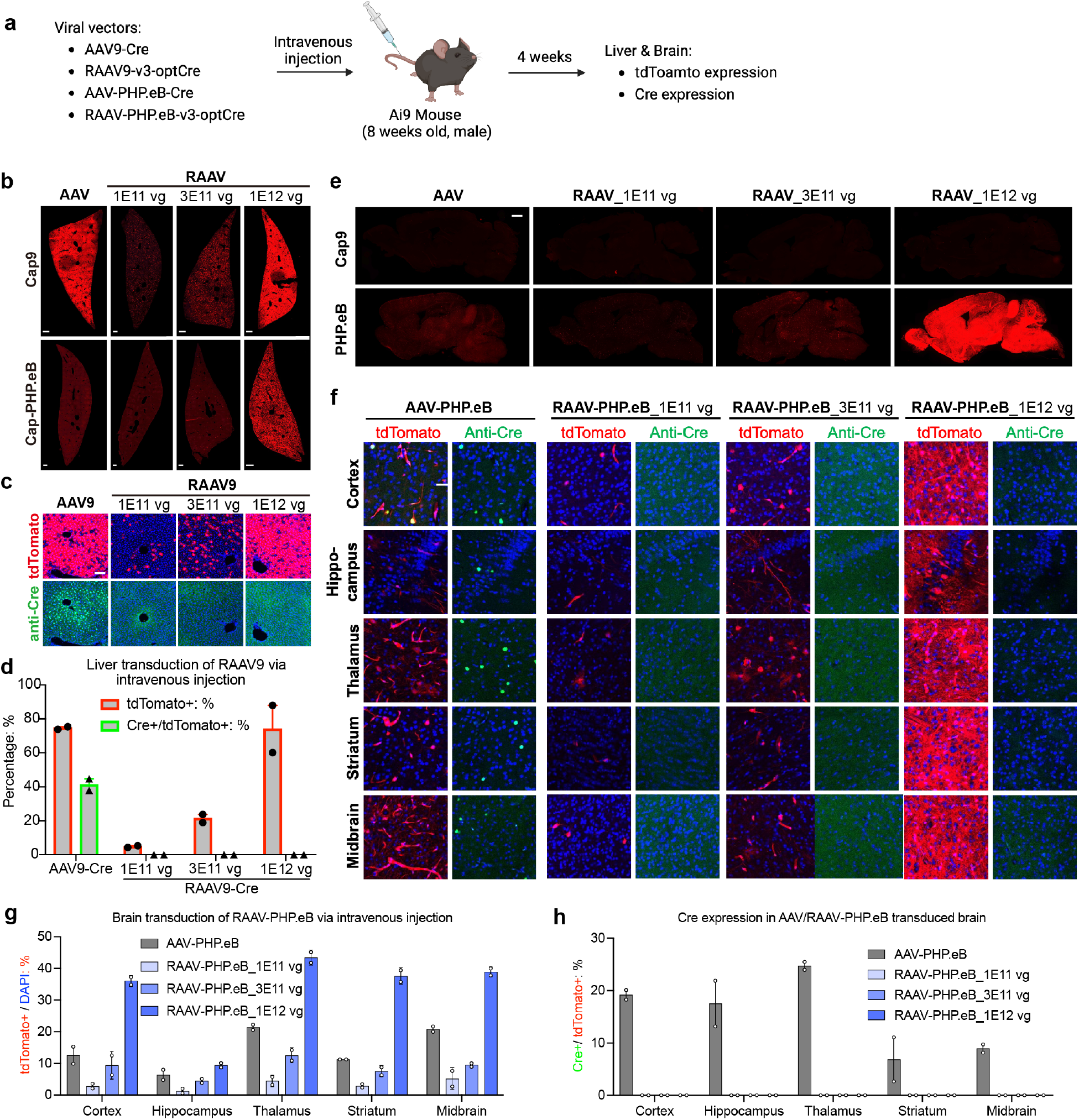
RAAVs enable tropism-dependent mRNA delivery to target tissues/organs to transiently express functional proteins. **a**, Schematic of RAAV *in vivo* systemic infection assay. RAAVs (carrying optCre mRNA) were generated by the RAAV-v3 system. As a control, AAVs carrying Cre DNA were generated and intravenously injected into Ai9 mice at a dose of 1E11 vg/mouse. **b**, Transduction of RAAVs/AAVs in the liver of Ai9 mice 4 weeks after intravenous injection. N=2 mice. Scale bars, 1000 μm. **c**, Fluorescence microscopy analysis of the expression of tdTomato and Cre in the liver of Ai9 mice 4 weeks after intravenous injection with RAAV9/AAV9. N=2 mice. Scale bars, 100 μm. **d**, Statistic analysis of tdTomato- and Cre-positive cells in (c). Data are shown as individual data points and mean values ± range for n = 2 mice. **e**, Transduction of RAAVs/AAVs in the brain of Ai9 mice 4 weeks after intravenous injection. N=2 mice. Scale bars, 1000 μm. **f**, Fluorescence microscopy analysis of the expression of tdTomato and Cre in the brain of Ai9 mice 4 weeks after intravenous injection with RAAV/AAV-PHP.eB. N=2 mice. Scale bars, 50 μm. **g** and **h**, Statistic analysis of tdTomato-(**g**) and Cre-(**h**) positive cells in (**f**). Data are shown as individual data points and mean values ± range for n = 2 mice.

In summary, by multi-step engineering, we have developed a mRNA delivery vector RAAV from the DNA virus AAV that exhibited high selectivity of mRNA and very low residual DNA packaging (∼ 0.005%). The RAAV systems combined the transient nature of mRNA with a variety of tissue tropism of the AAV capsid, making them ideal for either broad-spectrum or tissue-specific mRNA delivery. The results from our study demonstrate the feasibility of using rational engineering to change the virus genome type. Our strategy in developing RAAVs could be extended to other DNA viruses, in order to endow them the mRNA packaging capability. Further optimization of our RAAV system could be achieved by improving the specificity of RNA packaging, vector productivity, vector quality, as well as the integrity, stability and translation efficiency of the packaged genome, especially for large-size RNAs. RAAV-mediated gene editing exhibits a lower off-target editing rate compared to AAV-mediated methods (Fig. 4g,h), and the transient expression of Cas in the absence of DNA may offer potential advantages of reduced genome integration and reduced immune responses to Cas proteins, suggesting a safer approach for *in vivo* gene editing (Fig. 4i). Although RAAV may has a similar packaging capacity to conventional AAV, the emergence of compact RNA-guided nucleases with comparable editing efficiency to Cas9 ^39^, makes them suitable for delivery via a single RAAV. Nevertheless, our RAAV represents the first BBB-crossing mRNA delivery system that could efficiently infect the whole brain at therapeutic doses, a distinctive capability not yet achieved by existing RNA delivery tools, making it ideal for basic neuroscience studies and therapeutic applications.

## Acknowledgments

We thank M. Poo for helpful discussions and comments on this manuscript, Y. Kong and L. Pan (Electron Microscopy Facilities of Center for Excellence in Brain Science and Technology, Chinese Academy of Science) for assistance with EM sample preparation and EM image analysis, L. Li and W. Wu for assistance in RNAscope assay, and H. Tang for NGS data analysis.

## Author contributions

W.B. and L.S. conceived the project. W.B. and H.Y. designed the experiments, T.L., Y.Y. and X.Y. assisted in the experimental design. W.B., D.Y., Y.Z. (Yaqing Zhao), G.L., Z.L., P.X. H.Q., X.W. (Xiaoqing Wu), P.C. and X.W. (Xuchen Wang) performed experiments, X.K. and H.Z. assisted with experiments. W.B., D.Y., Y.Z. (Yaqing Zhao), Z.L., G.L., X.K. and Y.Z. (Yingsi Zhou) analyzed the data. H.Y. and L.S. supervised the research and experimental design. W.B. wrote the manuscript draft and W.B. and H.Y. contributed to the editing of the manuscript.

