## Supplementary information for "Multi-step engineered adeno-associated virus enables whole-brain mRNA delivery"

### Methods

#### Cell culture

Human embryonic kidney cells (HEK-293T), mouse embryonic fibroblasts (MEFs) and Hela cells were maintained at 37°C with 5% CO2 in DMEM (Hyclone, H30243.01) supplemented with 10% fetal bovine serum (Gibco, 10099-141C), 1% MEM Non-Essential Amino Acids Solution (Gibco, 11140050) and 1% Penicillin-Streptomycin-Glutamine (Gibco, 10378016).

#### Plasmids

Plasmids were cloned using PCR amplification with Phanta Max Super-Fidelity DNA Polymerase (Vazyme, P505-d1) and assembled with NEBuilder HiFi DNA Assembly Master Mix (NEB, E2621L).

#### Mice

Homozygous Ai9 mice were obtained from the Jackson Laboratory. Heterozygous Ai9 mice were derived from crossing wild-type C57BL/6J females with homozygous Ai9 males. All housing and procedures were performed according to protocols approved by the Institutional Animal Care and Use Committees (IACUC) of HUIDAGENE Therapeutics Co., Ltd. All mice were housed in a room maintained on a 12 h light and dark cycle with ad libitum access to standard rodent diet and water. Animals were randomly assigned to various experimental groups. The AAVs and RAAVs were injected into the hippocampus by stereotaxic injection, and into the mice by intravenous injection.

#### Production of AAVs and RAAVs

Both AAVs and RAAVs were produced and purified in an identical manner. HEK293T cells were maintained in DMEM with 10% fetal bovine serum in 150-mm dishes and passaged every 2-3 days. Cells were seeded at 1.5E7 cells per 15 cm dish one day before polyethyleneimine (Polysciences 24765-1) transfection. Then, 15 μg AAV/RAAV transgene plasmid, 15 μg AAV/RAAV packaging plasmid and 30 μg pAd-Helper were transfected per plate. The day after transfection, the media was exchanged for fresh DMEM with 2% fetal bovine serum. The supernatants of transfected cells were collected on day 2 and day 5 post transfection. Cells were also scraped with a rubber cell scraper on day 5, pelleted by centrifugation for 10 min at 3000 g, resuspended in 500 μL hypertonic lysis buffer per plate (10 mM Tris base, 150 mM NaCl and 10 mM MgCl2) and lysed via three repeated cycles of freeze/thaw. Add 125 U $mL^{-1}$ Benzonase nuclease (Sigma, E1014-25KU) to the cell lysate and incubate at 37°C for 1 h to remove cellular nucleic acids and residual plasmids. The collected supernatants were mixed with a 5× solution of 40% poly (ethylene glycol) (PEG) in 2.5 M NaCl (final concentration: 8% PEG/500 mM NaCl), incubated on ice overnight to facilitate PEG precipitation, and spun at 3000 g for 15 min. The pellet was resuspended in 500 μL lysis buffer per plate and also treated with 100 U $mL^{-1}$ Benzonase nuclease (Sigma, E1014-25KU) at 37°C for 1 h. Combine the resuspended virus from concentrated supernatants with cell lysates, and the obtained crude virus were clarified by centrifugation at 3000 g for 10 min and added to Beckman Quick-Seal tubes (Beckman, 342414) via Cotton-plugged Sterile Pasteur Pipets (Kimble, 63B95P). A discontinuous iodixanol gradient was formed by sequentially floating layers: 9 mL 15% iodixanol in lysis buffer with 1 M NaCl, 7 mL each of 25 and 40% iodixanol in lysis buffer, and 5 mL 58% iodixanol in lysis buffer. Phenol red at a final concentration of 1 μg $mL^{-1}$ was added to the 25 and 58% layers to facilitate identification. Ultracentrifugation was performed using a Type 70 Ti rotor in an OPTIMA XE-90 Ultracentrifuge (Beckman Coulter) at 68,000 rpm for 1 h 30 min at 18°C. Following ultracentrifugation, 5 mL of solution was withdrawn from the 40–58% iodixanol interface via a 14-gauge needle, dialyzed with PBS containing 0.001% F-68 using 100-kD MWCO columns (EMD Millipore). The concentrated viral solution was sterile-filtered using a 0.22-μm filter. The final AAV/RAAV preparation was aliquoted and stored at -80°C until use.

**Extraction and quantification of viral genome**

The purified AAVs and RAAVs were first subjected to nuclease treatment at 37°C for 3 hours to remove unencapsidated DNA and RNA. After digestion with nucleases, the encapsidated genome of AAVs and RAAVs was extracted using the previously described method for extracting capsid-associated DNA[40]. The extracted viral genomes were directly subjected to qPCR and rt-qPCR to quantify the viral DNA titer and RNA titer. Pairs of primers were designed targeting AAV/RAAV genomes.

**Transmission electron microscopy analysis**

In sample preparation for negative stain-electron microscopy, 10 μL purified AAVs and RAAVs were dropped on a 300 meshes copper grid coated with a continuous carbon film. The sample was allowed to adsorb for 2 minutes after which excess solution was removed with kimwipes. Subsequently, 10 μL negative staining solution containing 3% aqueous phosphotungstic acid was dropped on the TEM grid and incubated for 2 minutes, the excess solution was removed by touching the edge with kimwipes. The sample was allowed to dry before observation under a Talos L120C transmission electron microscope with a magnification of 73,000. Images were taken using Thermo Scientific™ CETA 16 4Kx4K CMOS camera.

**Silver stain**

Samples from purified AAV/RAAV vectors were loaded onto 4-20% Bis-Tris Gradient Precast Gels (Tanon, 180-9115H) and ran using 1xMOPS running buffer (Tanon, BT8100-2002). The gels were stained with Fast Silver Stain Kit (Beyotime, P0017S).

**Analysis of viral genome on agarose denaturing gel**

Mix 10 ng viral genomes with a 0.5 volume of Glyoxal Load Dye (Invitrogen, AM8551), and incubate the samples at 50°C for 1h. The denatured viral genomes were subsequently separated on a 1% glyoxal denaturing agarose gel (added 1/5000 SYBR™ Green II) at room temperature for 1h. The image was taken by the Tanon 2500 series automatic gel image analysis system.

**Mouse embryonic fibroblasts (MEFs) isolation**

Embryos from homozygous Ai9 mice were isolated between E12.5 and E18.5. After the heads, tails, limbs, and most of the internal organs were removed, the embryos were minced and trypsinized for 20 min and then seeded into 10 cm cell culture dishes in 10 mL of complete DMEM media. The cells were split at 1:2–1:3 ratios when freshly confluent, passaged two or three times to obtain a morphologically homogenous culture, and then frozen or expanded for further studies.

**HEK293T Cre reporter cell line generation**

HEK293T Cre reporter cell line was generated with the PiggyBac transposon system. A PB-T-loxP-tdTomato cassette was generated by subcloning the loxP-tdTomato cassette from Ai9 (Addgene #22799) into a PB-T plasmid. HEK293T reporter cell lines were created by seeding cells at 50% confluency in 6 well plates. The following day, the PB-T-loxP-tdTomato construct were cotransfected with the helper plasmid pCAG-PBase using polyethylenimine. Transfected cells were selected in puromycin (Thermo Fisher, A1113803) for 2 weeks and then sorted based on GFP on a BD FACSAria™ III Cell Sorter. Single sorted-cells were deposited into 96-well plates to get monoclonal cell lines.

**mRNA-sequencing of whole cell RNA and viral vector genomes**

RAAV and its no-MCP control were generated (with ten 15cm-dishes per group) and VLP RNA was extracted as viral vector genome extraction described above. Whole cell RNA was extracted with TRIzol Reagent (Invitrogen, 15596018) and purified using the phenol-chloroform extraction method. Subsequently, 1 μg RNA was used for the following library preparation. To mitigate potential bias resulting from the small amount of VLP RNA used during library preparation, we supplemented the viral vector genomes with 1 μg of carrier RNA. The poly(A) mRNA isolation was performed using Oligo(dT) beads, and the multiplexed RNA sequencing library was prepared using VAHTS® Universal V8 RNA-seq Library Prep Kit for Illumina (Vazyme, NR605). Libraries were sequenced on the Illumina novaseq 6000 using a 2x150 paired end (PE) configuration according to the manufacturer's instructions. Quality control was performed using Cutadapt (V1.9.1, phred cutoff: 20, error rate: 0.1, adapter overlap: 1bp, min. length: 75, proportion of N: 0.1). Clean data were aligned to reference genome (GRCh38.p13 + optCre) via software Hisat2 (v2.2.1). Differential gene expression analysis was performed using the DESeq2 Bioconductor package, a model based on the negative binomial distribution. Estimation of dispersion and logarithmic fold changes incorporate data-driven prior distributions, with Padj of genes set to<=0.05 to detect differentially expressed ones. Full read alignments were generated using Geneious prime.

**AAV and RAAV infection**

For all Ai9-MEFs infection experiments, cells were plated on 48-well plates at a density of 5E4 cells per well 24 hours before infection. Purified AAVs and RAAVs were added to Ai9-MEFs in triplicate. Vector genome titer was used for MOI calculation. Infected cells were collected at different time points for analyzing Cre DNA, Cre RNA, and Cre protein, or maintained for 5 days before flow cytometry analysis. To investigate the source of viral RNAs in infected cells, the transcription inhibitor - actinomycin D (AAT Bioquest 17505) was added to the cells at a concentration of 5 μg/mL 2 hours post-infection.

For HEK293T Cre reporter cell line infection experiments, cells were plated on 48-well plates at a density of 8E4 cells per well 24 hours before infection. Purified AAVs and RAAVs were added to HEK293T Cre reporter cells in triplicate. Vector genome titer was used for MOI calculation. Cells were maintained for 5 days before flow cytometry analysis.

**qPCR and RT-qPCR**

Total cellular DNA was extracted with TIANamp genomic DNA kit (TIANGEN, DP304-03). Total cellular RNA was extracted with TRIzol Reagent (Invitrogen, 15596018) and purified using the phenol-chloroform extraction method. Total RNA was reverse transcribed using the HiScript II Q RT SuperMix for qPCR (+gDNA wiper) (Vazyme, R223-01) according to the manufacturer's guidelines. qPCR was performed using AceQ qPCR SYBR Green Master Mix (Vazyme, Q111-02) on a CFX96 Touch™ Real-time PCR System (Bio-Rad) according to manufacturer's guidelines.

**Flow cytometry analysis**
Five days after transduction, transduced Ai9-MEFs or HEK293T Cre reporter cells were washed once with 1x PBS and dissociated with 0.25% trypsin-EDTA. Cells were resuspended with DMEM (containing 10% FBS), and the rescued tdTomato signals were determined using flow cytometry (Beckman CytoFlex). Analysis was performed using FlowJo v10.7 (BD Biosciences). Representative gating schemes are shown in Fig. S5.

**RNAScope assay**
HeLa (human cervical carcinoma) cells were seeded in 8-well glass chamber slides (MERCK, #PEZGS0816) at a density of 8E3 cells per chamber 24 hours before infection. Cells were then infected with RAAV-DJ (MOI=10,000 vg) or AAV-DJ (MOI=1,000 vg) in DMEM (containing 2% FBS). Bafilomycin A1 (Selleck, #S1413) was applied to the cells 1 h prior to infection at a concentration of 100 nM and kept in the medium for 24 h, concomitantly with the infection. At 1 or 6 h after transfection, actinomycin D (AAT Bioquest, #17505) was added at the final concentration of 5 µg/ml. At various time points post-infection, the cells were fixed and processed for RNAscope analysis.

RNAscope assay was performed according to the manufacturer's protocols of RNAscope™ Multiplex Fluorescent Reagent Kit v2 (ACD, #323100). Briefly, fixed cells were pretreated using the Universal Pretreatment Reagents (ACD, #322380). The chemically modified Cre probe (ACD, #474001) consists of 22 ZZ pairs. The pretreated cells were hybridized with the target probes at 40°C for 2 h and labeled with TSA Vivid fluorescent dye 520 (ACD, #323271) at a concentration of 1:1000. Nuclei are visualized using DAPI staining. The imaging was performed using a confocal microscope (Nikon C2si, Nikon).

**Western Blot**
For all the western blotting experiments, cells were lysed in LDS Sample Buffer (Biofuraw 180-8201D). The proteins were separated using SDS–polyacrylamide gel electrophoresis and transferred to polyvinylidene difluoride membranes. The membranes were blocked by 5% fat-free milk dissolved in TBS/0.05% Tween-20 (TBST) for 1 h, and incubated with anti-Cre monoclonal antibodies (1:1000, Cell Signaling Technology, 15036S) at 4 °C for 3 hours, washed 5 times in 1x TBST, incubated with anti-rabbit secondary antibodies (1:1000, Cell Signaling Technology, 7074S) for 1 h at room temperature, washed 3times in 1x TBST, then imaged with Tanon 4600. Tubulin detected using anti-tubulin polyclonal antibodies (1:3000, Bioworld, AP0064).

**Stereotaxic injection (into hippocampus) & Intravenous injection**
To investigate the infectivity of AAVs and RAAVs in mice hippocampus, Ai9 Mice (8 weeks old) were anesthetized with a mixture of zoletil (60 µg/g) and xylazine (10 µg/g), and then unilatrally, stereotactically injected with 1 µL AAV-Cre (two doses were set: 1E8 vg/mouse and 1E7 vg/mouse) or 1 µL RAAV-Cre (1E8 vg/mouse) into the dentate gyrus region of the right hippocampus according to the following coordinates: anteroposterior (A/P) = -1.7 mm, mediolateral (M/L) = -1.0 mm, dorsoventral (D/V) = -2.1 mm.

To investigate the tropisms of AAVs and RAAVs in mice brain, Ai9 Mice (8 weeks old) were anesthetized and intravenously injected with 300 µL AAV-Cre (1E11 vg/mouse) or RAAV-Cre (three doses were set: 1E11 vg/mouse, 3E11 vg/mouse, and 1E12 vg/mouse).

**Immunofluorescence staining and imaging of tissues.**
To investigate the infectivity and persistence of AAVs and RAAVs in mice, paraformaldehyde-fixed cryostat tissue section samples (brain and liver) were prepared 4 weeks after injection. Tissue sections were stained with anti-Cre antibodies (1: 800, cell signaling techonology, 15036S) and followed by Alexa Fluor 488-AffiniPure Donkey Anti-Rabbit IgG (H+L) (1: 1000, Jackson ImmunoResearch, 711-545-152). The nuclei were stained by DAPI (D3571, Invitrogen) and mounted with SlowFade Diamond Antifade Mountant (Invitrogen, S36972) on glass slides. The imaging was performed using a confocal microscope (Nikon C2si, Nikon).

**gRNA guide cell line generation**
A guide against human *TTR* gene was cloned using NEBuilder HiFi DNA Assembly under the control of a U6 promoter into a custom PB-T vector. HEK293T cells were seeded at 50% confluency in 6 well plates. The following day, the PB-T-U6-gRNA construct was cotransfected with the helper plasmid pCAG-PBase using polyethyleneimine. Transfected cells were selected in puromycin (Thermo Fisher, A1113803) for 2 weeks and then sorted based on BFP on a BD FACSAria$^{TM}$ III Cell Sorter. Single-sorted cells were deposited into 96-well plates to get monoclonal cell lines.

**Indel sequencing of *in vitro* edited cells**
*In vitro,* 96-well plates of tissue culture cells were infected with AAV-DJ-Cas12Max and RAAV-DJ-Cas12Max, and cells were lysed with 20 µL lysis buffer from One Step Mouse Genotyping Kit (Vazyme, PD101-01) 5 days after infection. The target region was amplified from genomic DNA by nested PCR. Barcoded PCR products were pooled together, purified with Gel Extraction Kit (OMEGA, D2500-02), and sequenced on an Illumina HiSeq system (150-bp paired-end reads). Indels were quantified from the resulting library using the script which has been deposited on github (https://github.com/yszhou2016/Cas12f/blob/main/0.Cas-Finder/3.Indel_Calculate.pl).

**Statistics**
Data were analyzed using GraphPad Prism 8. Quantitative data are presented as mean ± SD with n = 3 biological replicates per condition. Unless otherwise stated, biological replicates represent independent treatments in separate virus batches, culture wells or mice. Statistical significance was computed using unpaired t-test. The specific statistical method applied, and descriptions of replicates are provided in the figure legends. The asterisks indicate statistical significance; unless otherwise specified, *P < 0.05, **P < 0.01, ***P < 0.001, ****P < 0.0001; ns, non-significant.

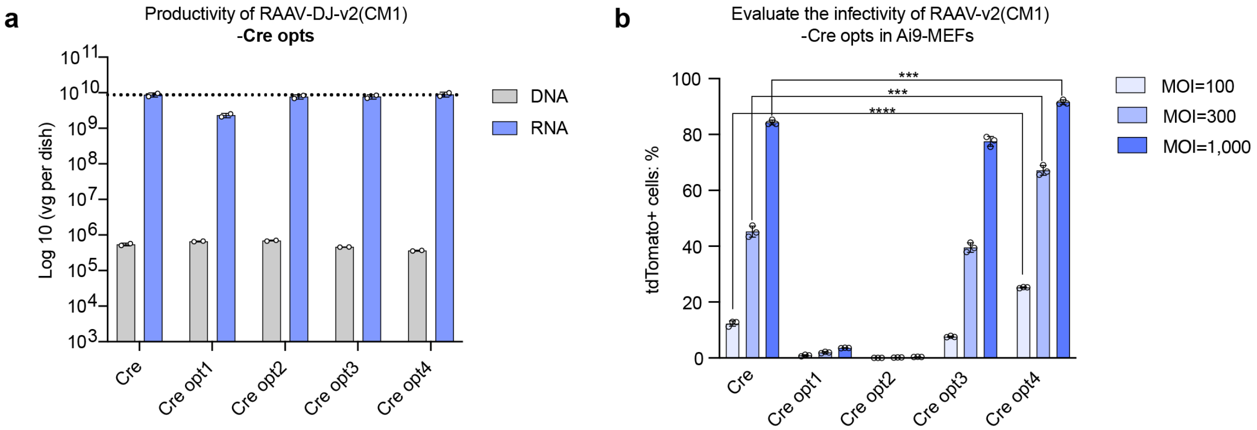


**Fig. S1 Cargo sequence optimization for improving RAAV infectivity. a**, Quantification of the encapsidation of different Cre coding sequences in RAAVs by qPCR and RT-qPCR using primers targeting WPRE. Cre opt sequences were obtained by codon optimization using several online tools. RAAVs were generated by the RAAV-v2 (harboring the combined mutation 1 in helicase) system. Data are shown as individual data points and mean values ± range for n=2 biological replicates. **b**, Investigating the infectivity of RAAVs carrying different Cre coding sequences by analyzing the percentage of tdTomato+ Ai9-MEFs 5 days after infection by cytometry. An artificial capsid DJ was used to produce AAV/RAAV vectors. Vector genome titer was used for the MOI calculation. Data are mean values ± SD with n = 3 biological replicates; unpaired, two-tailed t-test; ***P < 0.001, ****P < 0.0001.

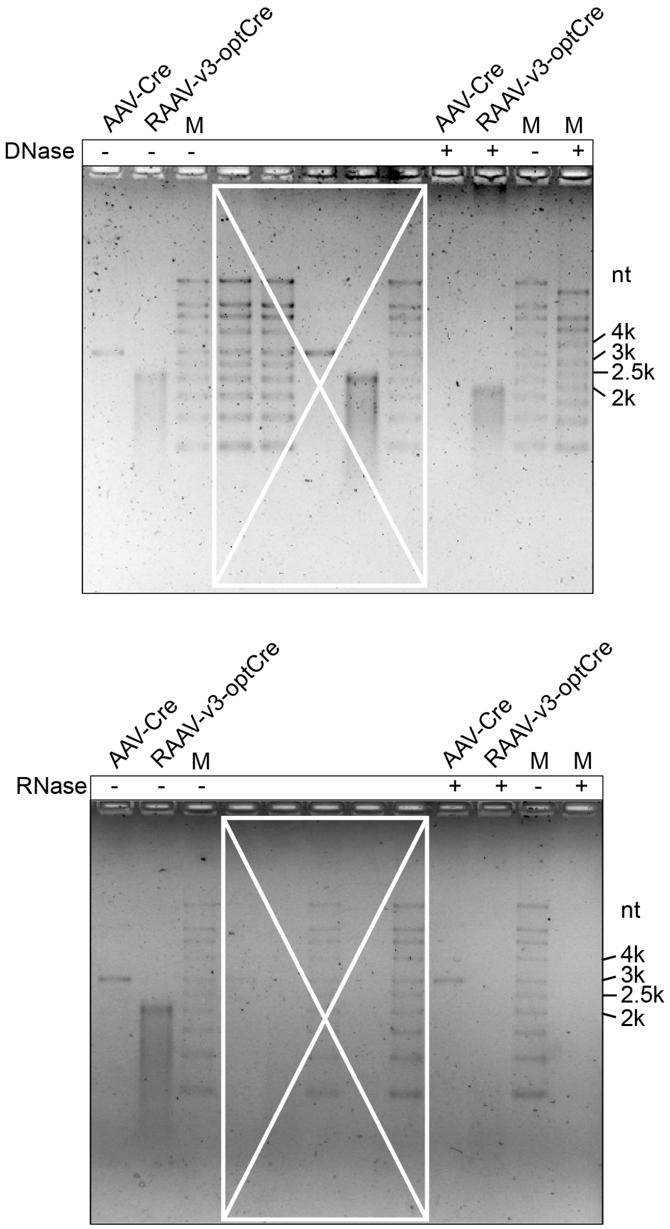


**Fig. S2 Analyzing RAAV genome on denaturing agarose gels stained with SYBR™ Green II.** DNaseI and RNaseI treatment groups were set to identify the RAAV genome. These are full images of Fig. 3d, the lanes within the white boxes are irrelevant samples.

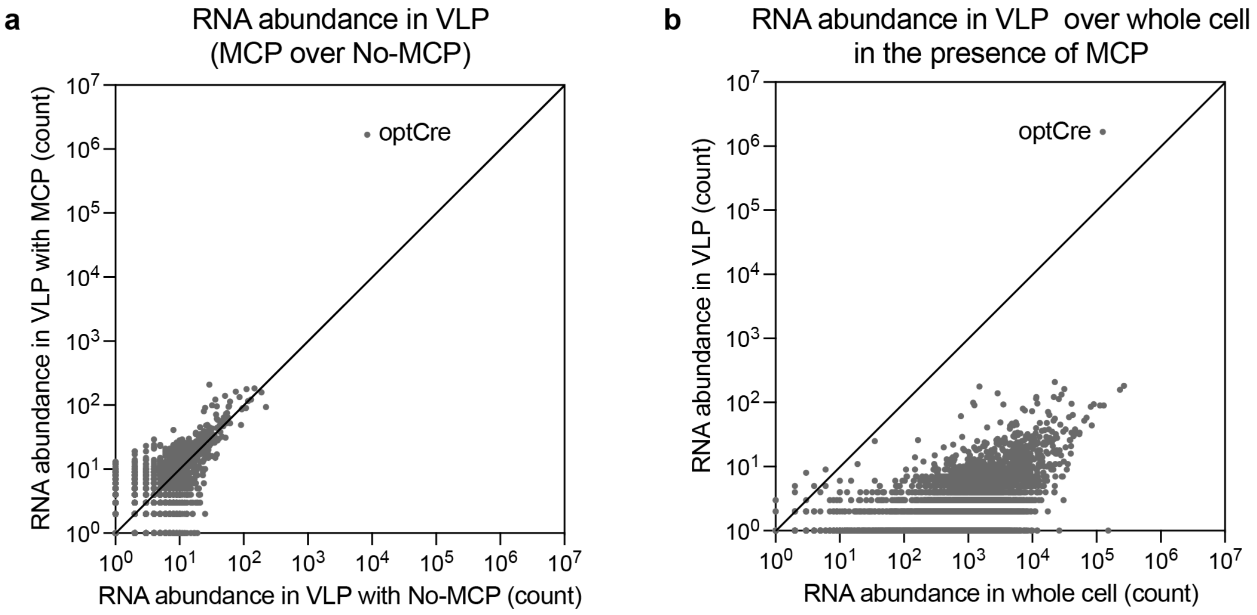


**Fig. S3 RAAV packaging specificity. a**, Differential RNA abundance of the VLP fraction in the presence or absence of MCP. **b**, Only RPS-harboring mRNA (optCre) was efficiently packaged in RAAV.

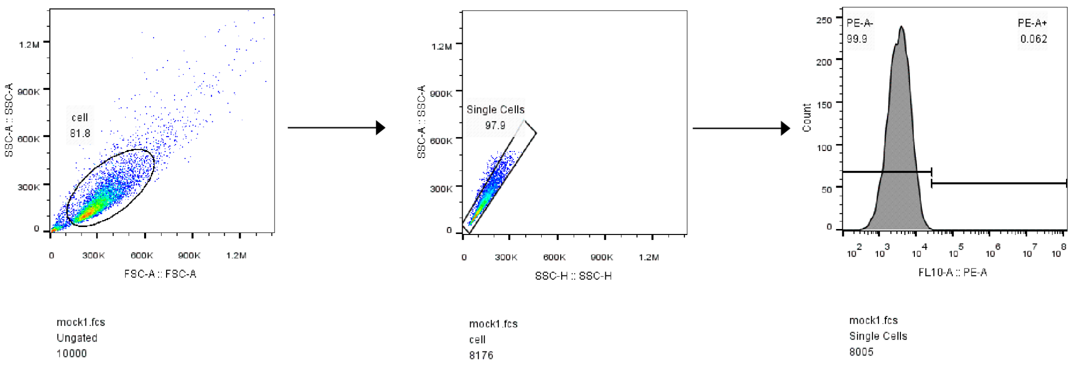


**Fig. S4 Representative flow cytometry gating scheme for RAAV/AAV *in vitro* Ai9-MEF infection experiments.** Cells were first gated on FSC and SSC to remove debris. Following the singlets were gated on SSC. The tdTomato+ cells were gated based on uninfected controls (mock).

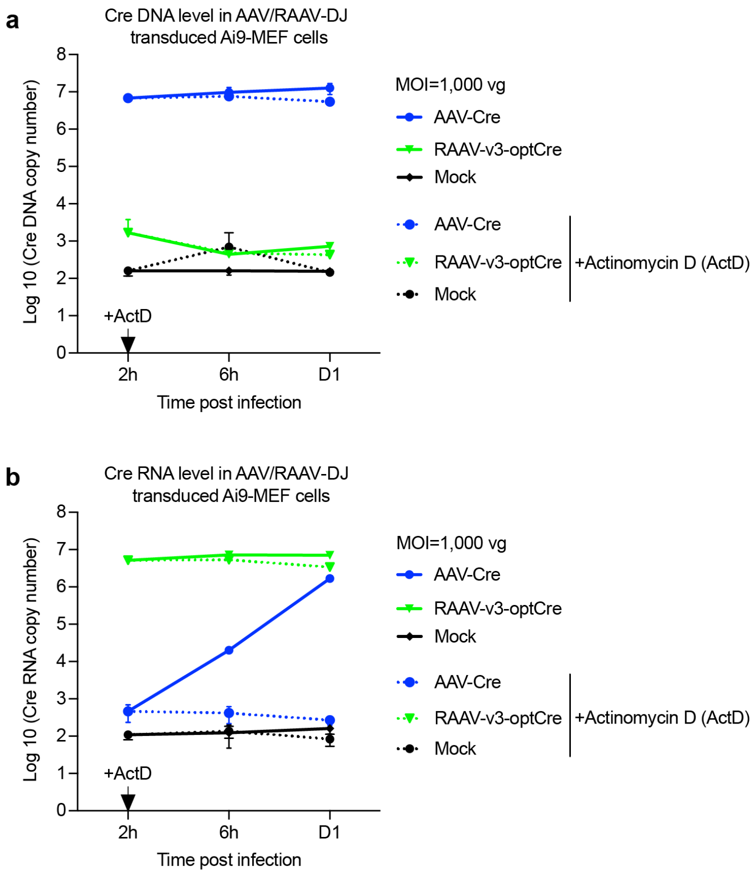


**Fig. S5 Investigate the effects of the transcription inhibitor actinomycin D on the viral DNA and RNA levels in AAV/RAAV-DJ-infected Ai9-MEFs. a,b,** Analyzing the viral DNA (**a**) and RNA (**b**) levels in AAV/RAAV infected Ai9-MEFs. Vector genome titer was used for the MOI calculation. Ai9-MEFs were infected by AAV-Cre or RAAV-v3-optCre at MOI 1000 vg, actinomycin D was added to the cells at a concentration of 5 μg/mL 2 hours post-infection, and cells were collected at 6- and 24-hours post-infection for analyzing viral DNA and RNA. Mock: unifected control. Data are shown as mean values ± SD for n = 3 biological replicates.

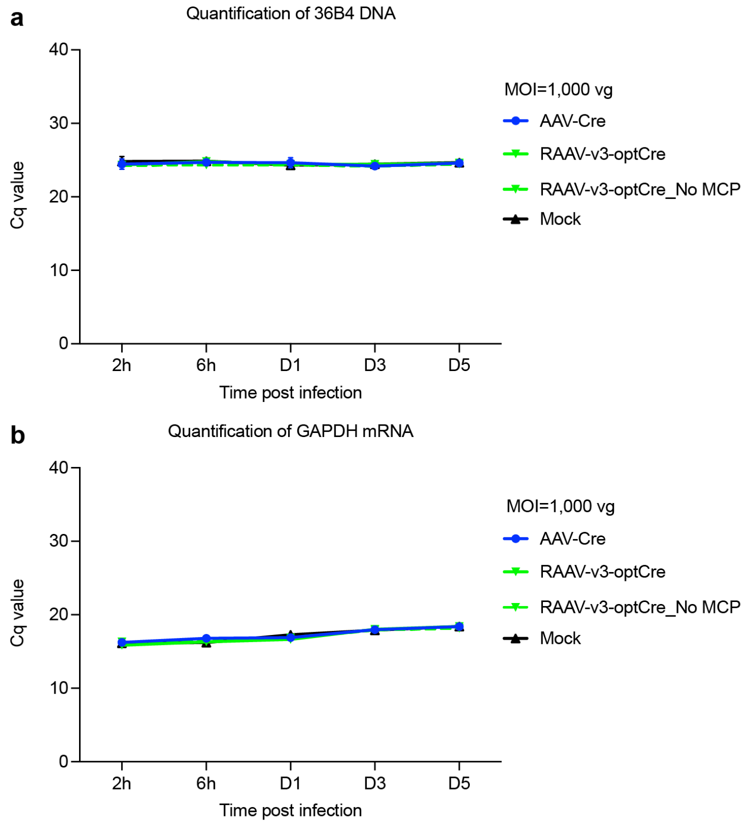


**Fig. S6 Cellular DNA/mRNA were analyzed as loading controls in AAV/RAAV-DJ infected Ai9-MEFs. a**, Ct value of cellular housekeeping gene (36B4). **b**, Ct value of cellular GAPDH mRNA. Mock: uninfected control. Data are shown as mean values ± SD for n = 3 biological replicates.

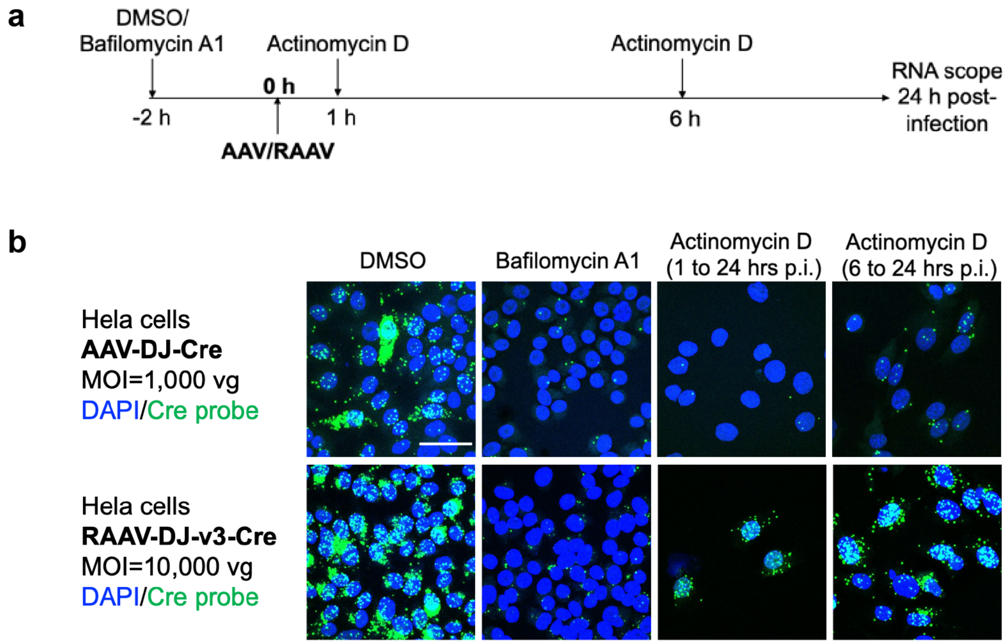


**Fig. S7 Investigate the effects of the vacuolar H+-ATPase inhibitor bafilomycin A1 and the transcription inhibitor actinomycin D on AAV/RAAV-DJ-mediated infection. a**, Schematic of the experiment. Hela cells were treated with bafilomycin A1 (100 nM) 2 h prior to infection with AAV-DJ-Cre (MOI 1,000 vg) or RAAV-DJ-v3-Cre (MOI 10,000 vg), and the transcription inhibitor actinomycin D was added at a concentration of 5 µg/mL 1 h or 6 h post-infection. DMSO was set as solvent control. At 24 hours post-infection, the cells were fixed and processed for RNAscope analysis. **b**, The effects of the vacuolar H+-ATPase inhibitor bafilomycin A1 and the transcription inhibitor actinomycin D on AAV-/RAAV-mediated transduction. Nuclei were visualized using DAPI staining. Viral RNA and DNA were detected with a DNA probe that binds to the Cre mRNA and DNA. Scale bars, 50 µm.

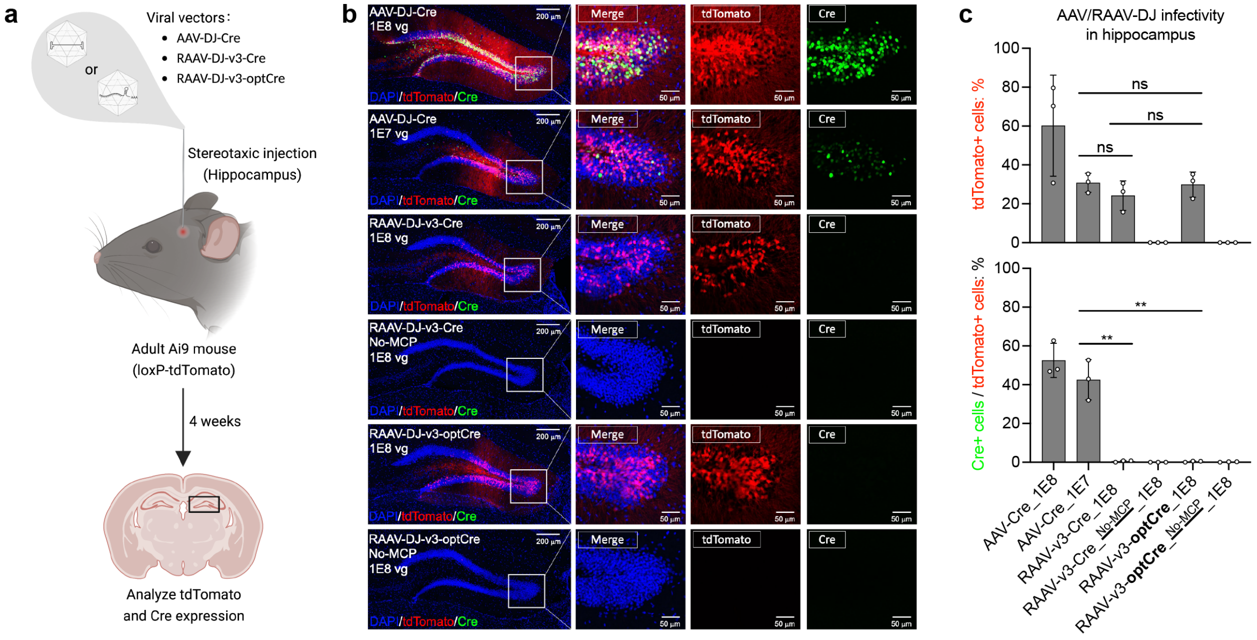


**Fig. S8 RAAVs enable efficient transfer of mRNA to the mouse hippocampus to transiently express functional proteins. a**, Experimental workflow for RAAV/AAV-DJ in vivo hippocampus transduction. DNA amounts were normalized between RAAV and RAAV with no MCP (negative control). **b**, Fluorescence microscopy analysis of the expression of tdTomato and Cre in hippocampi of Ai9 mice 4 weeks after stereotaxic injection with RAAV/AAV. The right images are magnified versions of the area indicated by the white box in the left images. Scale bars, 200 µm in left images and 50 µm in right images. **c**, Statistic analysis of tdTomato- and Cre-positive cells in (**b**). Data are shown as individual data points and mean values ± SD for n = 3 mice; unpaired, two-tailed t-test; $^{**}P < 0.01$; ns, not significant.

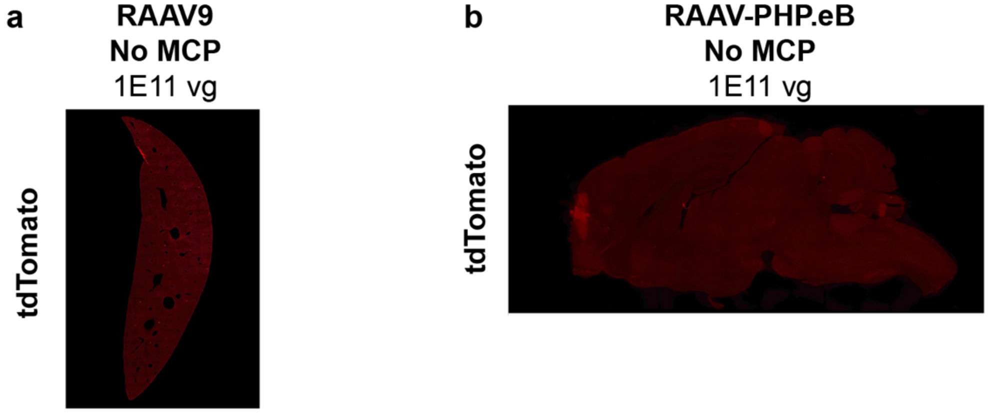


**Fig. S9 RAAV with No-MCP showed no infectivity. a**, Infection of RAAV9 with No-MCP in the liver of Ai9 mice 4 weeks after intravenous injection. N=2 mice. **b**, Transduction of RAAV-PHP.eB with No-MCP in the brain of Ai9 mice 4 weeks after intravenous injection. N=2 mice.

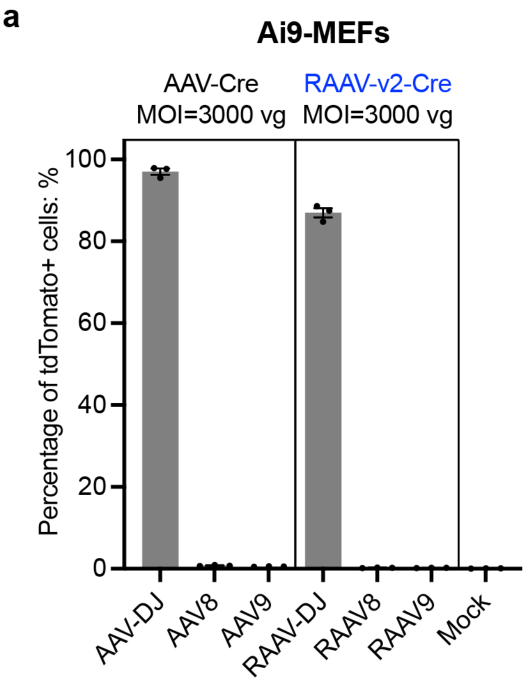


**Fig. S10 Investigating the cellular tropism of RAAVs. a,b**, Comparison of the infectivity of AAVs and RAAVs in Ai9-MEFs (**a**) and HEK293T Cre reporter cells (**b**). RAAVs were generated by RAAV-v2. Vector genome titer was used for the MOI calculation. Investigating the infectivity of AAV and RAAV by analyzing the percentage of tdTomato+ cells 5 days after infection by cytometry. Data are shown as individual data points and mean values ± SD for n = 3 biological replicates.
